# VapC12 ribonuclease toxin modulates host immune response during *Mycobacterium tuberculosis* infection

**DOI:** 10.1101/2023.08.24.554572

**Authors:** Shaifali Tyagi, Srikanth Sadhu, Taruna Sharma, Abhijit Paul, Manitosh Pandey, Vaibhav Nain, Deepak Rathore, Samrat Chatterjee, Amit Awasthi, Amit Kumar Pandey

**Author notes:** Correspondence: Amit Awasthi Amit Kumar Pandey.

## Abstract

Mechanistic understanding of antibiotic persistence is a prerequisite in controlling the emergence of MDR cases in Tuberculosis (TB). We have reported that the cholesterol-induced activation of VapC12 ribonuclease is critical for disease persistence in TB. In this study, we observed that relative to the wild type, mice infected with Δ*vapC12* induced a proinflammatory response, had a higher pathogen load, and responded better to the anti-TB treatment. In a high-dose infection model, all the mice infected with Δ*vapC12* succumbed early to the disease. Finally, we reported that the above phenotype of Δ*vapC12* was dependent on the presence of the TLR4 receptor. Overall, the data suggest that the inability of Δ*vapC12* to resolve neutrophil-mediated inflammation reduced bacterial killing by altering the T-cell response. In conclusion, our findings suggest the role of the VapC12 toxin in modulating the host’s innate immune response in ways that favor the long-term survival of the pathogen inside the host.

## Introduction

COVID-19 has augmented the tuberculosis (TB) burden by disrupting the existing healthcare systems resulting in a delay in diagnosis and treatment [1]. In spite of the availability of anti-tubercular therapy, its eradication still remains obscure [2]. In addition, non-compliance to a prolonged treatment regimen fuels the development of multidrug and extensively drug-resistant TB posing blockades to global TB control [3]. The virulence of *Mycobacterium tuberculosis (M. tuberculosis*) is highly transmissible primarily through the aerosolization of the respiratory secretions from the infected individuals. This together with a relatively low virulence has allowed this pathogen to successfully co-evolve with humans for centuries without causing any symptoms or death [4]. We believe that the complex network of host-pathogen interaction, thus developed during evolution, defines TB in terms of its presentation either as an asymptomatic infection or an acute infection presented with clear symptoms. Nevertheless, both the pathogen and the host have gradually evolved mechanisms that would identify and neutralize any mutually directed threats.

Innate immunity is an indispensable arm of the host immune system mediated by phagocytes such as macrophages; bona fide host cells harboring the mycobacteria [5]. The role of neutrophils which are highly motile innate immune cells has been shown in experimental models of TB pathogenesis [6–9]. Following the *M. tuberculosis* infection, these are the primary cells to percolate the lungs and have an extremely intricate role in TB pathology [10]. Neutrophils mediate their phagocytic activity by producing proteases, reactive oxygen and nitrogen intermediates and antimicrobial peptides [11]. These immune cells are the most common cell type present in the sputum, bronchoalveolar fluid and cavities of active TB patients [10, 12]. There is a negative correlation between their abundance in the blood of *M. tuberculosis-*infected patients and disease severity [13–16]. On the contrary, some neutrophil subpopulations are also known to aid in providing a permissive replicating niche and nutrient reservoir for *M. tuberculosis*, indicating their role in promoting mycobacterial persistence [17, 18]. The same has also been reported in *M. tuberculosis* infected rhesus macaques [19] and zebrafish [20]. Growing evidence suggests the phenotypic and functional diversities of neutrophils which extend well beyond their conventional role as antimicrobial agent. Timely resolution of neutrophils at the inflamed site dictates their fate as either protective or dysregulated immune cells and is crucial for the effective restoration of the excessive inflammatory response to homeostasis. The ineffective clearance of neutrophils leads to exacerbation of inflammatory response and perpetuation of tissue damage [21]. Furthermore, the role of mycobacterial genes in shaping an appropriate neutrophil-mediated innate immune response and its timely resolution, which further affects the downstream adaptive immunity, remains largely unknown.

Previously, using the guinea pig model, we demonstrated that both lungs and spleen isolated from guinea pigs infected with *M. tuberculosis* lacking the VapC12 ribonuclease toxin were presented with higher bacterial burden and increased lung pathology relative to the wild type (WT) [22]. We hypothesize that the VapC12 toxin in *M. tuberculosis* is critical for modulating the host immune response and hence the absence of this toxin in *M. tuberculosis* resulted in a severe form of the disease. To further our understanding of the role of the host immune system in imparting the above phenotype to the *vapC12* deleted strain, in the current study, we performed comprehensive immune-profiling studies in mice using the WT, Δ*vapC12* and the complemented strains to identify host correlates of disease persistence. We found that in comparison to WT, the lungs of mice infected with the mutant strain were presented with higher bacterial load and enhanced pathology. Additionally, these mice showed an excess of myeloid cell infiltration (neutrophils and myeloid-derived suppressor cells [MDSCs]) in the lungs with an alleviated T-cell immune response. In the survival study, performed with a high-dose infection model, while all the mice infected with WT strain survived none of the mice from the Δ*vapC12* group survived 18 weeks post-infection. Additionally, lung transcriptome analysis revealed that the observed neutrophil accumulation was mirrored by increased *S100A8* and *S100A9* mRNA levels in the lungs infected with Δ*vapC12* when compared with the WT strain. Interestingly, we found that the enhanced pathogenesis observed in the mice infected with Δ*vapC12* was dependent on the presence of TLR-4 receptors. These results suggest that vapC12 ribonuclease mediated regulation of expression of pathogen-associated molecular patterns (PAMPs) in *M. tuberculosis* modulates host immune response which is essential for the long-term survival of the pathogen inside the host. A better understanding of the mechanism of generation and maintenance of antibiotic persistence in the context of host-pathogen interaction will go a long way in designing novel intervention strategies against tuberculosis. Host-directed therapies (HDT) targeting such pathways in combination with the existing anti-TB treatment can increase the efficacy of the current TB regimen.

## Methods

### Bacterial strains and culture conditions

All mycobacterial strains were grown and maintained in Middlebrook 7H9 broth and 7H11 agar (Difco Middlebrook 7H9 broth [catalog no. 271310] and 7H11 agar [catalog no. 283810]) and supplemented with 10% OADS (bovine serum albumin, oleic acid, and dextrose); 0.05% Tween was added in 7H9 broth to enhance bacterial growth. Bacterial enumeration by CFU was examined on 7H11 agar plates. The concentrations of antibiotics used were as follows: hygromycin B (50 µg/mL) and kanamycin (25 µg/mL).

### Mouse infection study

Animal experiment protocols were reviewed and approved by the animal ethics committee of Translational Health Science and Technology Institute India [(IAEC/THSTI/183), (IAEC/THSTI/167)], Faridabad, India. Animal experiments were performed in accordance with guidelines provided by the Committee for the Purpose of Control and Supervision of Experiments on Animals (Govt. of India). Pathogen-free C57BL/6 and C3H/HeJ mice were obtained from Small Animal Facility, THSTI. As described earlier, briefly, prior to infection, mid-log phase cultures (O.D =0.8-1) of H37Rv (WT), Δ*vapC12* and Δ*vapC12:vapBC12 M. tuberculosis* strains were harvested, washed and single-cell suspensions were prepared[23]. Six to eight-week old mice were infected by aerosol exposure in the Glas-Col airborne infection apparatus with approximately 50-100 CFUs of different strains. Mice from each group (N = 5) were sacrificed at different time points as indicated in the experiments (Day 0, 2, 4 and 8 weeks post infection). The bacillary load was determined by removing lungs and spleen of infected mice and the serial dilution of the organ homogenates was plated on 7H11 agar plates supplemented with 10% OADS for bacterial enumeration. The lung sections were treated with 10% formalin and hematoxylin and eosin stained were used for histopathological analysis. A pathologist who was unaware of the samples’ identities evaluated the coded tissue samples for granulomatous organization. The pathologist scored all the granulomas in each section and added them up to calculate the total granuloma score. For drug clearance study, after establishing bacterial infection till week 4, mice were treated with 10mg/kg RIF through oral gavaging till week 8. Also, a separate untreated group was kept as a control. Bacillary enumeration was done on week 4 and 8 by aseptically removing organs and homogenizing them. The homogenized organs were then serially diluted and plated on 7H11 agar (containing Polymyxin A, amphotericin B, vancomycin, trimethoprim, carbenicillin and cycloheximide mix).

### Flow Cytometry Analysis

Mice were sacrificed at 2 and 4 week post-infection, and their lungs, spleen and LN (draining lymph node) were isolated. Single-cell suspension was prepared and stained (1 x 10^6^ cells) with fluorescently labelled anti-mouse CD11b (eBioscience Invitrogen, USA), anti-mouse Gr1 (eBioscience Invitrogen, USA), anti-mouse F4/80-FITC (eBioscience Invitrogen, USA), anti-mouse CD68 (eBioscience Invitrogen, USA), anti-mouse CD206 (eBioscience Invitrogen, USA), anti-mouse CD80 (eBioscience Invitrogen, USA), anti-mouse NK1.1 (eBioscience Invitrogen, USA), anti-mouse CD3 (BD Biosciences USA), anti-mouse CD4 (BD Biosciences USA), anti-mouse CD8 (eBioscience Invitrogen, USA) and anti-mouse γδ-TCR (eBioscience Invitrogen, USA). Briefly, for surface staining, cells were stained for 20 minutes at room temperature, and were washed twice with PBS. Stained samples were then fixed for 20 minutes using 4% paraformaldehyde. Data was acquired on FACS Canto (BD Biosciences) and analyzed using FlowJo software (Tree Star, Ashland, OR, USA) [24, 25].

### RNA Sequencing

For transcriptome analysis, lung tissues of mice infected with different strains (4-week infected WT, Δ*vapC12* and Δ*vapC12:vapBC12*) were homogenized by bead beating, and total RNA was isolated using RNeasy mini kit according to the manufacturer’s protocol (Qiagen catalog no. 74104). RNA was eluted in nuclease-free water, subjected to DNA removal using TURBO DNase. NEB Ultra II directional RNA-sequencing Library Prep kit protocol was used to prepare libraries for total RNA-Seq (NEB, catalog no. E7760L). An initial concentration of 500ng of total RNA was taken for the assay. mRNA molecules are captured using magnetic Poly(T) beads (NEB, catalog no. E7490L) following purification, the enriched mRNA was fragmented using divalent cations under elevated temperatures. The cleaved RNA fragments were copied into first-strand cDNA using reverse transcriptase. Second strand cDNA synthesis was performed using DNA Polymerase I and RNase H enzyme. The cDNA fragments were then subjected to a series of enzymatic steps that repair the ends, tails the 3’ end with a single ‘A’ base, followed by ligation of the adapters. The adapter-ligated products were then purified and enriched using the following thermal conditions: initial denaturation 98°C for 30sec; 12 cycles of - 98°C for 10sec, 65°C for 75sec; final extension of 65°C for 5mins. PCR products are then purified and checked for fragment size distribution on Fragment Analyzer using HS NGS Fragment Kit (1-6000bp) (Agilent, catalog no. DNF-474-1000). Genes with low expression value (mean FPKM value<0.5) were treated as background (i.e. ‘noise’) [26] and hence excluded from further analysis. After the noise removal, the gene expression values for each experimental condition were merged, and the final data comprised the FPKM value of 13796 genes. To assess the variation in the gene expression data, we performed the principal component analysis (PCA) using the *prcomp* package in R, which applies the singular value decomposition to the data matrix. For the comparison between gene expression profiles from mice infected lungs with WT and Δ*vapC12* strain of *M. tuberculosis*, genes with a fold change of 2 and false discovery rate (FDR)-adjusted p-value <0.05 were considered differentially expressed. The Benjamin-Hochberg multiple hypothesis testing [27] was applied to obtain the FDR-adjusted p-value. Subsequently, the analyses were performed on the MATLAB and R software. The *mafdr* function in MATLAB was applied to obtain the FDR-adjusted p-value. PCA plot, volcano plot and dot plot were generated in R using *ggplot2*, *enrichplot* and *gplots* packages. Heatmaps were built using heatmap.2. Pathway enrichment analysis was performed in R using the *clusterProfiler* package.

### Quantitative RT-PCR

Total RNAs from lungs were isolated using RNeasy Micro kit (Qiagen), and the cDNAs were synthesized by reverse transcriptase (RT) using the high capacity cDNA archive random priming kit (Applied Biosystems) according to the manufacturer’s recommended protocol. Real-time PCR was performed with 2 × SYBR mix (TaKaRa) in a QuantStudio-6 Flex Real-Time PCR System (Applied Biosystems) using the default run program. The relative expression of the target genes was calculated using the target threshold cycle value (Ct) and the 2^−ΔΔ*Ct*^ method with GAPDH as the internal loading control [28]. The primer sequences used for real time PCR are listed in **Table S1**.

### Depletion of Monocyte Derived Suppressor Cells

MDSCs were depleted using 200μg anti-mouse Ly6G (Bio X Cell) or isotype IgG (Bio X Cell). Briefly, in acute infections (less than 21 dpi), mice were given 200μg of anti-mouse Ly6G intra-peritoneally every other day between 10 and 20 dpi.

### Statistical Analysis

Each experiment was repeated at least three times independently and three biological replicates were used. Data were plotted as mean ± SEM. Student’s t-test (Mann-Whitney), two-way ANOVA and one-way ANOVA (Kruskal-Wallis) were performed wherever applicable by using Graph Pad Prism 7.0 software. For the survival curve, Kaplan Meier statistical analysis was performed. P-value < 0.05 was considered to be statistically significant.

## Results

### VapC12 ribonuclease toxin modulates growth, virulence and drug tolerance in *M. tuberculosis*

To replicate our earlier findings of the study performed on guinea pigs [22], we infected C57BL/6 mice with WT, *vapC12* null mutant and complemented *M. tuberculosis* strains. We found a significantly higher bacterial load in the lungs and spleen of mice infected with *vapC12* null strain as compared to the WT and complemented strains both at 4 and 8 weeks post-infection **(Figure 1A and 1B)**. In comparison to the WT, gross examination of the lungs infected with the mutant strain revealed higher inflammation and enhanced tissue damage. Histological examination of the lungs infected with the WT strain at 8 weeks post-infection revealed a typical necrotic granuloma in the centre of the lung (40X) with epithelioid cells and lymphocytes in the granuloma (100X) **(Figure 1C and Supplementary** Figure 1A). In contrast, at a similar time point post infection, lung tissues of mice infected with the *vapC12* null strain displayed a significantly higher number of granulomas **(Figure 1D)**. Mice infected with complemented strain had a phenotype similar to those infected with WT strain **(Figure 1C & 1D)**. Previously, using kill-curve experiments, we have demonstrated that the absence of the *vapC12* gene in *M. tuberculosis* reduced the frequency of generation of drug-tolerant population in presence of rifampicin under *in-vitro* growth conditions[22]. To replicate the above findings under disease conditions, we performed a drug clearance study using C57BL/6 mice infection model. We infected a group of mice with either WT and Δ*vapC12* strain and estimated the rate of pathogen clearance by estimating bacterial load in infected organs at different time points **(Figure 1E)**. Similar to previously reported guinea pig data, 4 week post infection, the organs infected with Δ*vapC12* had high bacterial load in comparison with the organs isolated from the WT infected mice (Figure 1F **&** 1G). Importantly, upon treatment with RIF and INH, the rate of decline in the bacterial load was higher in mice infected with Δ*vapC12* strain relative to WT at 8 weeks post infection (Figure 1F **&** 1G). To account for the differences in the initial bacterial load (week 0), we had to normalize the data relative to week 0 between H37Rv and *vapC12* mutant strains **(Figure 1 F & 1G).** Importantly, upon treatment with RIF we observed a ∼2 fold and ∼7 fold increase in the killing of the pathogen in the lungs and spleen respectively in the mice infected with Δ*vapC12* as compared to mice infected with the H37Rv **(Figure 1F & 1G**) whereas treatment with INH resulted in ∼10 fold better killing in the lungs and spleen of the mice infected with Δ*vapC12* relative of the H37Rv. These results show that the absence of *vapC12* gene in *M. tuberculosis* decreased the frequency of generation of drug-tolerant population potentiating the efficiency of RIF and INH treatment in infected mice **(Figure 1F & 1G)**. Since *vapC12* deleted strain of *M. tuberculosis* demonstrated enhanced pathogenicity, we measured the estimated time of survival of WT and *ΔvapC12* infected mice in a high dose infection model (500-1000 bacteria/lung) **(Supplementary** Figure 1B). Interestingly, we found that 16 weeks post-infection, all the mice (n=10) infected with the WT strain survived while none of the mice infected with the Δ*vapC12* survive (n=10) survived **(Figure 1H)**. Collectively, these data suggested that absence of *vapC12* imparted hypervirulence phenotype and reduces the frequency of generation of persisters in mice infected with *M. tuberculosis*.

**Figure 1:**
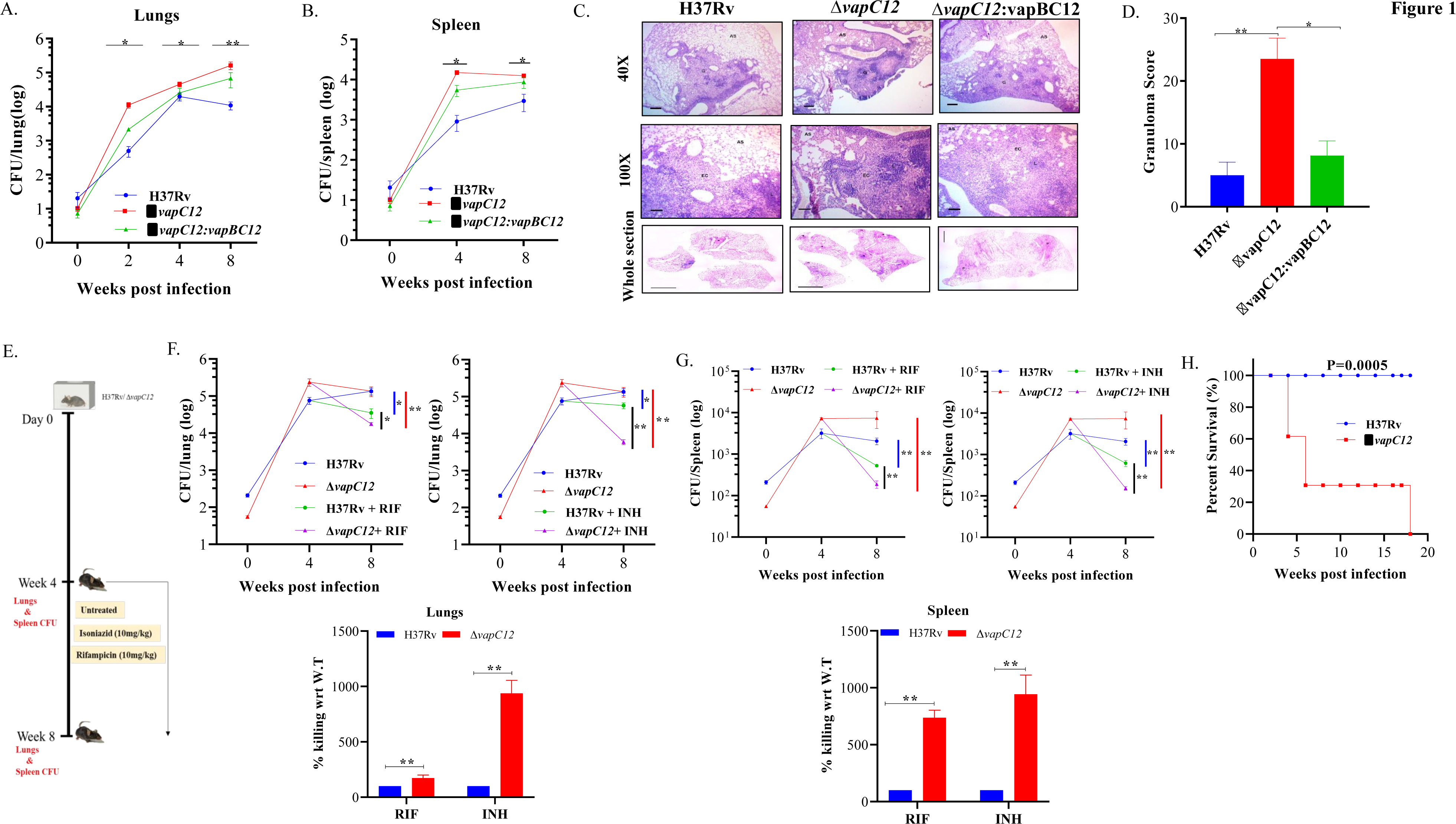
*vapC12* mutant strain of *M. tuberculosis* modulates growth and drug tolerance. (A and B) Bacterial load in the (A) lungs or (B) spleen of mice infected with H37Rv, *ΔvapC12*, and *ΔvapC12*:*vapBC12* strains of *M. tuberculosis* at designated time points, the lungs were homogenized in 2 ml of saline, and ten-fold serial dilutions of homogenates were plated on 7H11+OADC plates. Each group constituted five mice per time point. Data plotted represent the mean ± SEM. Significant differences observed between groups are indicated. Data were analyzed using the Mann–Whitney test with **P < 0.01 and *P <0.05 (C) Photomicrographs of H&E-stained (40× and 100×) and whole section image of lung sections from mice infected with different strains of *M. tuberculosis* at 8 weeks post infection. G=Granuloma, As=Alveolar Space, EC=Epithelioid Cells and L=Lymphocytes (D) Granuloma fraction of the lung tissue samples of mice infected with different strains of *M. tuberculosis*, based on the semi-quantitative estimation of the fraction of the lung tissue covered with granuloma. Data were analyzed using the Mann–Whitney test with *P < 0.05 and **P < 0.01. (E) Schematic representation of mice infection experiment. Mice (n = 5 per group at each time point) were infected with H37Rv and Δ*vapC12*. After the establishment of infection for 4 weeks, antibiotics (INH or RIF) were orally administered to the mice for the next 4 weeks. A parallel group of mice was left untreated (control). Bacterial loads were enumerated in the lungs (F) and spleens (G) at different time points post-infection as depicted by scatter plot. Each data point represents the log10 CFU of an infected animal in individual organs, and the error bar depicts the SEM for each group. Bar graph represents the percent killing of the *vapC12* mutant strain with respect to WT at week 8 post-infection in the (F) lungs and (G) spleen. Data was normalized with the untreated group and plotted with respect to H37Rv (100%) on the y-axis. Data were analyzed using the Mann–Whitney test with **P < 0.01 and *P <0.05. (H) Percent survival plot of mice (n=20) infected with H37Rv and *ΔvapC12* strains for 18 weeks post infection. Data were analyzed using the log-rank test with ***P value 0.0005.

### VapC12 toxin modulates host immune response during *M. tuberculosis* infection

To understand the *vapC12* mediated differences in the host immune response, we performed detailed immune-profiling studies on organs isolated from mice infected with WT, Δ*vapC12* and complemented **s**trains of *M. tuberculosis* at week 4 post-infection **(Figure 2A)**. We estimated the innate immune cell population; monocytes, macrophages, MDSCs, neutrophils, and Natural Killer-T (NKT) cells in lungs and spleen isolated from infected mice **(Figure 2B**, **2C and Supplementary** Figure 2A). A set of uninfected mice was included as a control group in all the experiments. We found that relative to the WT, both lungs and spleen isolated from mice infected with Δ*vapC12* had a significantly lower percentage (∼2%) of monocytes population **(Figure 2B & 2C)**. In contrast, relative to the WT, organs isolated from the mice infected with the *vapC12* mutant strain had a significantly higher percentages of macrophage, neutrophil and MDSCs **(Figure 2B & 2C)**. The restoration of the mutant specific phenotype to the WT levels in the group of mice infected with the complemented strain further confirms the role of VapC12 toxin in modulating the host immunity. As reported in other settings, accumulation of MDSCs parallels disease progression in lung parenchyma and exacerbate disease by suppressing T-cell functions [29–32] . Our findings corroborated the previous studies as the MDSCs population in the lungs and spleen was significantly elevated by ∼3-fold in the mutant strain in comparison to the WT infected group **(Figure 2B & 2C)**. Neutrophil infiltration in the lungs; which is a common feature of disease severity in TB patients [10] and susceptible mice [18, 33, 34] was found to be 3-fold higher in the *vapC12* mutant strain as compared to the WT and complemented strain **(Figure 2B & 2C)**. Unlike lungs, we did not observe any differences in the neutrophil population in the spleen isolated from either the WT and *vapC12* infected mice, suggesting that the neutrophil-mediated inflammation by the *vapC12* mutant was specific to the lungs **(Figure 2C)**. It has been reported that individuals with pulmonary TB have lower number of NKT cells as compared to the healthy controls, indicating that they may have a protective role in TB infection [35, 36]. Surprisingly, we found that the percentage of NKT cells in lungs and spleen isolated from *vapC12* mutant infected mice were 2-fold and 3-fold elevated as compared to the organs isolated from WT infected mice **(Figure 2D)**. Together, these data indicated that *vapC12* mutant mediates immunosuppression by accumulating MDSCs and promoting neutrophil infiltration in the lungs.

**Figure 2:**
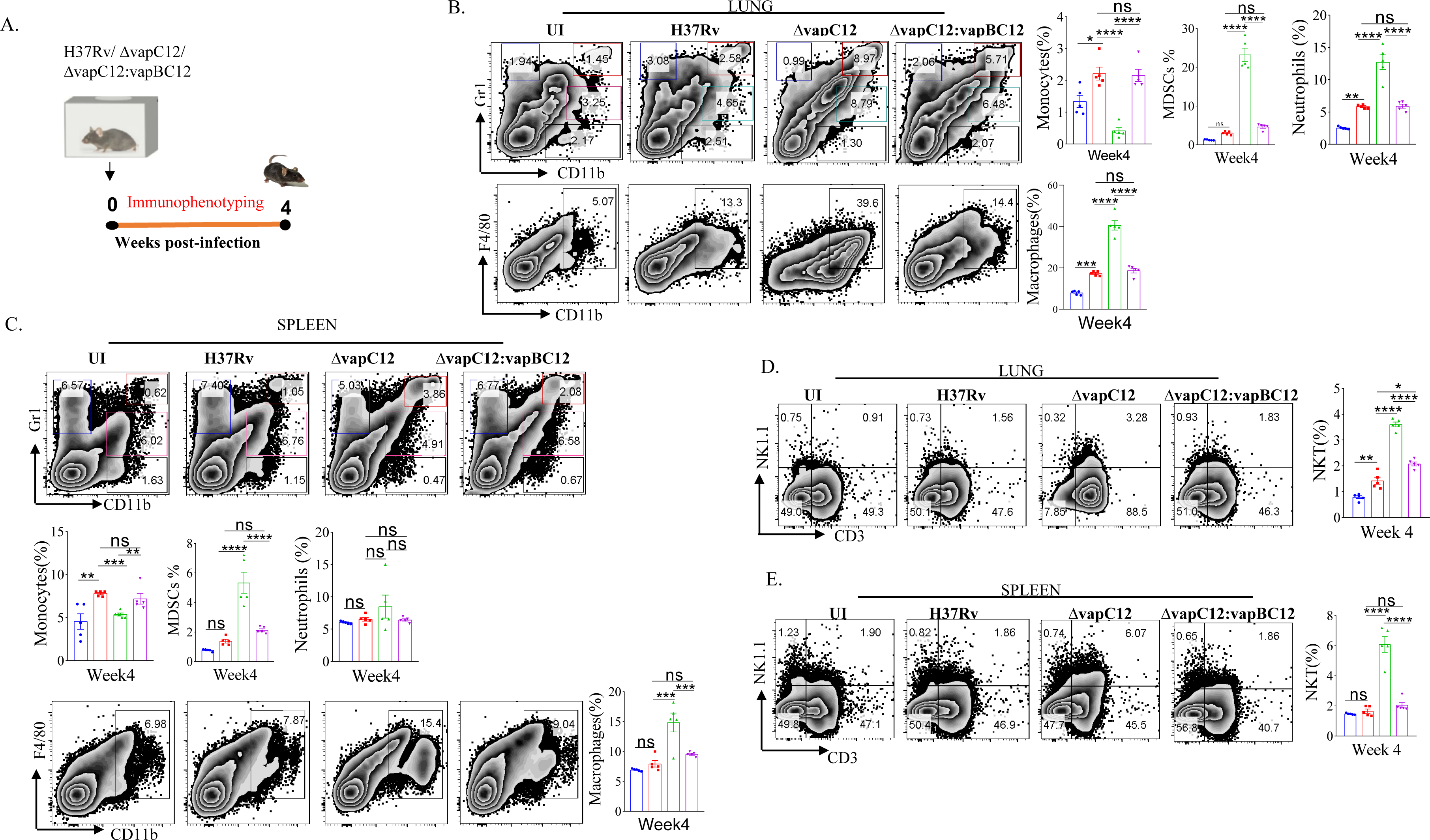
VapC12 toxin modulates host immune response during *M. tuberculosis* infection. (A) Schematic representation of the murine infection experiment. Mice (*n* =5 per group at each time point) were infected with H37Rv, *ΔvapC12*, and *ΔvapC12*:*vapBC12*. Immuno-profiling of lung infiltrated cells and splenocytes harvested at 4 weeks post infection. (B and C) Zebra plots and its representative bar graphs indicating the percentage frequency of monocytes (Gr1+), macrophages (CD11b+F4/80+), MDSCs (CD11b+ Gr1+) and neutrophils (CD11b+Gr-1^int^) cells harvested from (B) lungs and (C) spleen. Data are shown as mean ± SEM (n=5 animals per group). Data were analyzed using the one-way ANOVA followed by Tukey’s multiple comparison test (*P <0.05, **P < 0.01, ***P<0.0005, ****P<0.0001). (D and E) Zebra plots and its representative bar graphs showing the percentage frequency of NKT (NK1.1+CD3+) cells harvested from (D) lungs and (E) spleen. Data are shown as mean ± SEM (n=5 animals per group). Data were analyzed using the one-way ANOVA followed by Tukey’s multiple comparison test (*P <0.05, **P < 0.01, ***P<0.0005, ****P<0.0001).

### VapC12 toxin regulates the polarization of macrophages during *M. tuberculosis* infection

The choice of effector mechanism selected by the host to clear infection is regulated by the type of cytokine response dictated by the initial innate immune response. The fate of Th1 and Th2 polarisation, ideal for intracellular and extracellular pathogens respectively, is effectively decided by the initial differentiation of the macrophages to either classical or alternate types. As an effective host immune escape strategy, the pathogen tries to manipulate the macrophage differentiation process in its own favor by subverting the host cell metabolic pathways [37]. To investigate role of the *vapC12* gene in manipulating macrophage differentiation we used antibodies against CD80 and CD206, classical markers for M1 and M2 respectively to quantify the relative M1/M2 population in the lungs of mice infected with different *M. tuberculosis* strains. We found a ∼1.5-fold decrease in the frequency of M1 (CD80+) macrophages in lungs isolated from mice infected with *vapC12* mutant as compared to the WT. On the contrary, we found higher frequency of M2 expressing macrophages in the lung tissue infected with Δ*vapC12* **(Figure 3A and Supplementary** Figure 3A). The same was further validated by quantifying the transcripts of markers specific to either M1 or M2 macrophages. We estimated the transcripts levels of typical M1 [CD80 and CD86] and M2 markers [Arginase1 (Arg1), YM1 and Fizz1) by isolating RNA from the lungs of WT, *vapC12* mutant infected and uninfected mice. As expected, we found ∼6-fold (Fizz1), ∼10-fold (YM1) and ∼35-fold (Arg1) increase in the expression of M2 markers in the *vapC12* mutant strain as compared to the WT strain **(Figure 3B)**. In concordance with elevated levels of M2 markers, a clear upregulation in the gene expression levels of the Th2 related cytokines; IL 6 (****P<.0.0001) and IL10 (*P<.0.05) was also observed **(Figure 3C)**.

**Figure 3:**
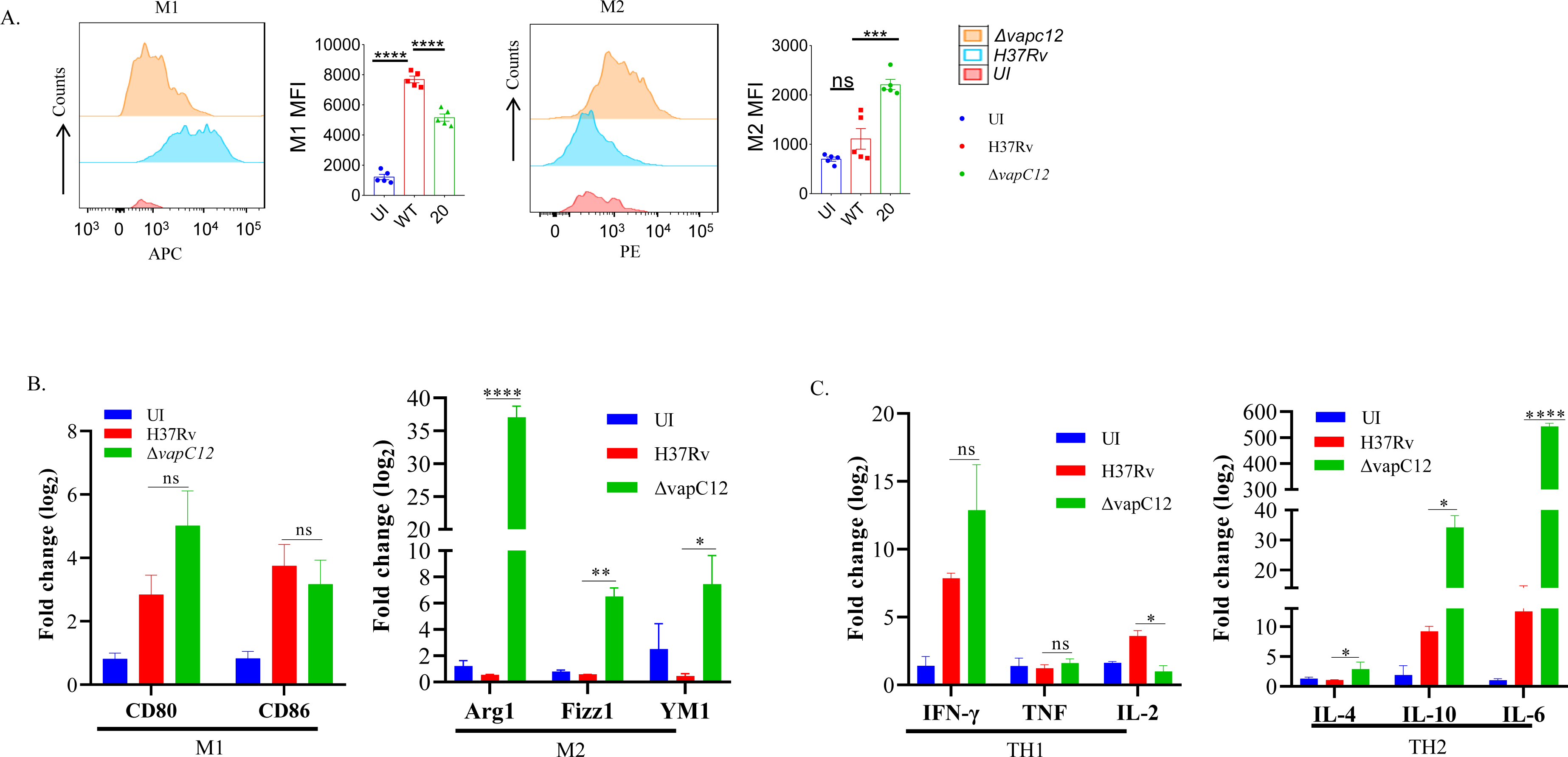
VapC12 toxin regulates the polarization of macrophages during *M. tuberculosis* infection. (A) Left panel M1: shows the histogram and bar graph of CD80 and right panel M2: shows the histogram and bar graph of CD206 in CD11b+ F4/80+ cells by flow cytometry and bar graph represents MFI (Mean fluorescent index). Data were analyzed using FlowJo software (tree star) and statistical calculation was presented as mean ± SEM, n = 5. ***P<0.0002, ****P < 0.0001. (B) Bar graph for mRNA expression of M1 (CD80 and CD86) and M2 (Arg1, YM1 and Fizz1) macrophage markers in the lung cells, data representative of triplicates *P < 0.05, **P < 0.01, ***P < 0.001, ****P < 0.0001 (C) Bar graph for mRNA expression of TH1 (IFN-γ, TNF and IL-2) and TH2 (IL-4, IL-10 and IL-6) in the lung cells, data representative of triplicates *P < 0.05, **P < 0.01, ***P < 0.001, ****P < 0.0001

Surprisingly, the transcription signature associated with M1 and Th1 markers CD80, CD86, IFN-γ and TNF-α showed non-significant changes in their gene expression levels and a clear reduction in Th1-induced secretion of IL-2 was noted **(Figure 3B and Figure 3C)**. Overall the data convincingly demonstrates that the ratio of M1/M2 (Th1/Th2) is significantly lower in the mice infected with Δ*vapC12* as compared to the WT strain. In sum, these data suggest that levels of VapC12 ribonuclease help *M. tuberculosis* control the M1/M2 polarization modulating the Th1/Th2 pathway critical for regulating the disease outcome.

### *vapC12* is critical for induction of adequate T cell immune response during *M. tuberculosis* infection

In addition to myeloid cells, CD4^+^ and CD8^+^ T lymphocytes are the key players in the establishment of protective immunity against TB in mice. The changes in cellular phenotype coincided with the significant decrease in the adaptive immune markers i.e. CD4^+^ and CD8^+^ T lymphocytes in the organs isolated from the *ΔvapC12* infected mice relative to the WT. In lungs harvested from mice infected with *vapC12* mutant strain, we observed a significant 3-fold and 2-fold reduction in the population of CD4^+^ and CD8^+^ T-cells respectively, both with p<0.05 as compared to the WT infected mice **(Figure 4A and Supplementary** Figure 4A). Surprisingly, while the CD4^+^ T-cell profile in the spleen for both groups were identical to the lungs, the CD8^+^ T-cell population remained unaltered **(Figure 4B)**. As shown in Figure 4C, both CD4^+^ and CD8^+^ T-cell populations were suppressed in the lymph nodes of *vapC12* infected mice. Interestingly, in comparison to the WT, the secondary lymphoid organs isolated from the mice infected with *ΔvapC12* harbored significantly higher numbers of γδ-T cell population 4 weeks post-infection **(Figure 4B and 4C)** with no such differences observed in the lungs **(Figure 4A)**. This data corroborates the previous findings which suggest the role of γδ-T cells in TB pathogenesis by inducing inflammation and granuloma formation [38]. Thus, we conclude that the absence of VapC12 toxin in *M. tuberculosis* prevents the induction of adequate T-cell response in the lungs and lymphoid organs (lymph nodes and spleen) of infected mice.

**Figure 4:**
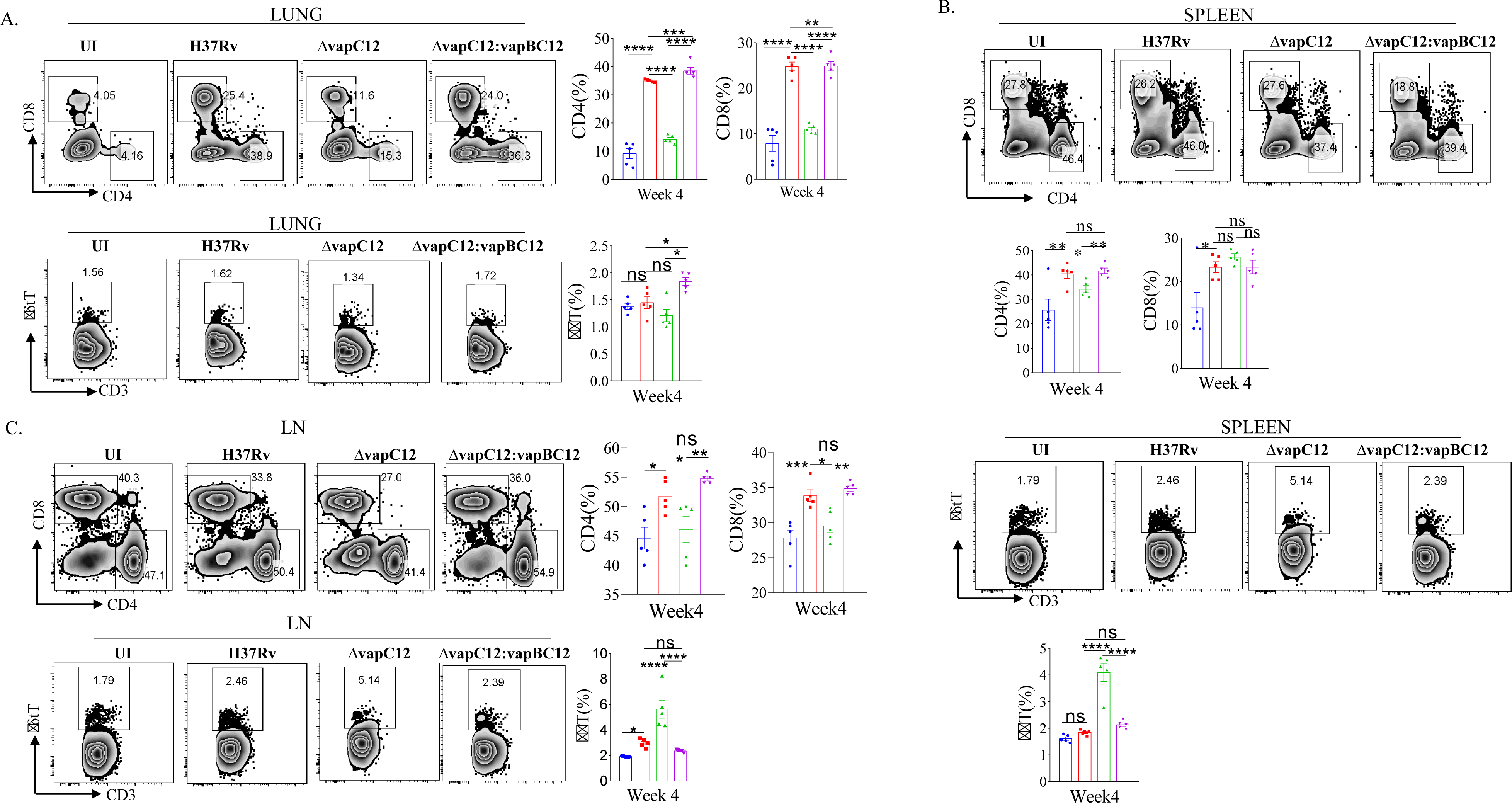
*vapC12* is critical for induction of adequate T cell immune response during *M. tuberculosis* infection. (A-C) Zebra plots and its representative bar graphs indicating percentage frequency of CD4, CD8 and γδ+T cells in the (A) lung (B) spleen and (C) Lymph nodes. Data were analyzed using FlowJo and shown as mean ± SEM (n=5 animals per group). Data were analyzed using the one-way Anova test followed by Tukey’s multiple comparison test (*P <0.05, **P < 0.01, ***p<0.0005, ****p<0.0001).

### VapC12 is essential for the resolution of neutrophil-mediated inflammation in tuberculosis

To further delineate the role of host factors associated with the hypervirulent phenotype of the mutant strain, we analysed the RNA sequencing data of host genes from lung tissues of mice infected with WT, Δ*vapC12* and complemented strain. The same was subjected to a detailed transcriptome analysis **(Figure 5A)**. Initially, a principal component analysis (PCA) was performed to visualize the variations in the obtained expression profiles. Data displayed in the first two principal components (PCs) suggest that all three gene clusters are well separated from each other (**Supplementary** Figure 5A). To study the VapC12-mediated differential expression of the host genes, we compared the gene expression profiles of lung tissues from mice infected with the Δ*vapC12* and the WT strains (**Figure 5B**). In total, 372 differentially expressed genes (DEGs) were obtained, out of which 82 and 290 genes were up- and down-regulated, respectively (**Supplementary** Figure 5B and **Supplementary File S1**). Pathway enrichment analysis of the upregulated genes revealed an enrichment in the processes related to neutrophil functions like neutrophil migration and neutrophil-mediated immunity **(Figure 5C)**. We found that in comparison to the WT, infection with the Δ*vapC12* strain enhanced the expression of genes that encode proteins implicated in neutrophil-mediated immunity in the host (Cxcl2, Trem1, S100A8/A9 and Ccl3) **(Figure 5D**). S100A8 and S100A9; cytosolic proteins of neutrophils are important for the recruitment of neutrophils to the granulomas as demonstrated in human and animal models of TB [39]. A ∼2.56 fold (log_10_) increase in the transcript levels of genes encoding *S100a8* and *S100a9* by the host infected with the *vapC12* mutant strongly suggests an increase in neutrophil recruitment and accumulation in the lung tissues. Genes belonging to the inflammation-associated markers like matrix metalloproteases (Mmp8 and Mmp9), secreted by neutrophils that are known to be critical for the formation of granuloma and inducing inflammation during the acute stage of infection [40], were found to be significantly induced in the lung tissues of mice infected with the mutant strain. We also found an increase in the expression of *Cxcr2* (22-fold)*, Cxcl1* (18-fold) and *Cxcl2* (177-fold) genes which suggests enhanced neutrophil recruitment in lungs of mice infected with *vapC12* mutant relative to the WT [41] **(Supplementary File S2)**. Additionally, IL1a (30-fold) and IL1b (19-fold), the pro-inflammatory cytokine markers associated with TB [42] were also found to be significantly upregulated in the tissue isolated from mice infected with mutant strain as compared to the WT and complemented strain **(Figure 5E**). Interestingly, in comparison to the WT, the expression of cystatins such as Cstdc4 [3.2-fold (log_10_)], Stfa2l1 [2.8-fold (log_10_)] and Csta2 [2.26-fold (log_10_)] which are the endogenous inhibitors of the host antimicrobial proteins cathepsins [43]were also increased in lungs of mutant strain infected animals **(Figure 5F**). Simultaneously, peptidases which play an important role in pathogen killing by activating the antimicrobial peptides were found to be significantly reduced in the *ΔvapC12* infected mice lungs **(Figure 5G & 5H**), setting off an unrestricted growth of *M. tuberculosis* leading to excessive inflammatory responses. Additionally, the observed reduction in the transcript level of genes associated with adaptive immune response correlates with our earlier findings that the *ΔvapC12* infected mice fail to induce an effective adaptive immune response **(Supplementary** Figure 5C). Together, these observations suggest that *ΔvapC12* proliferates and induces a proinflammatory response by modulating the transcript level of proteins that enhances neutrophil recruitment, IL-1 production, endopeptidase inhibitors and by suppressing the antimicrobial proteases and appropriate adaptive immune response inside the host.

**Figure 5:**
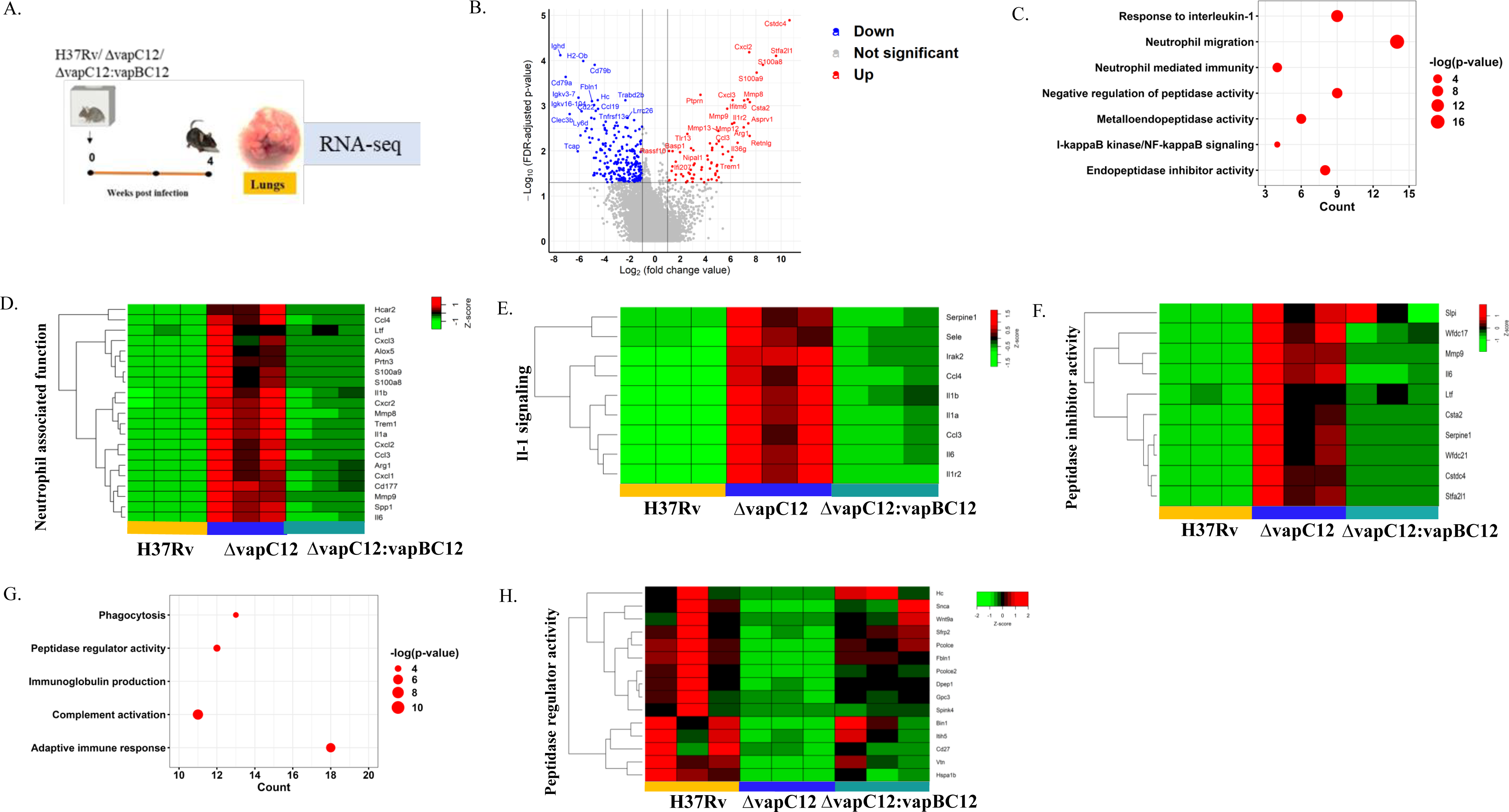
VapC12 is essential for the resolution of neutrophil-mediated inflammation in tuberculosis. (A) Pictorial representation of the process of animal experiment with mice. RNA isolated from lung tissues of mice (n=3) infected with WT, Δ*vapC12* and complemented strain after 4 weeks of infection. (B) The volcano plot showing comparison of gene expression profiles from mice infected with parental (WT) and Δ*vapC12* strain of *M. tuberculosis*. Differentially expressed genes (DEGs) were selected using the fold change cut-off value of 2 and FDR-adjusted p-value < 0.05. The up- and down-regulated genes are highlighted here in red and blue, respectively. (C) Dot plot of the gene ontology (GO) pathway enriched in upregulated genes. The x-axis represents the number of obtained upregulated genes for each pathway. (D) Heatmap representing the expression levels of the 21 upregulated genes involved in the neutrophil associated function. Data were Z-score transformed and the name of the genes are displayed on the right. The names of the corresponding infected strains for each sample are mentioned at the bottom of the figure. (E) Heatmap representing the expression levels of the 9 upregulated genes involved in the IL-1-mediated signaling. (F) Heatmap representing the expression levels of the 10 upregulated genes involved in the peptidase inhibitor activity. (G) Dot plot of the gene ontology (GO) pathway enriched in down-regulated genes. The x-axis represents the number of obtained upregulated genes for each pathway. (H) Heatmap representing the expression levels of the 15 downregulated genes involved in the peptidase regulator activity.

### Depletion of Myeloid-derived suppressor cells failed to restore the *vapC12* mutant phenotype

Apart from providing an intracellular environment conducive to the growth and the replication of *M. tuberculosis*, MDSCs are also known to evade adaptive immunity by suppressing T-cells responses [44–46]. As previously reported, MDSCs depletion can be achieved by the administration of 1A8 antibody [47]. To confirm the role of MDSCs in Δ*vapC12* induced hypervirulent phenotype we infected groups of C57BL/6 mice, with different *M. tuberculosis* strains, followed by treatment with either 1A8 antibody or their isotype control **(Figure 6A)**. Treatment with 1A8 antibody resulted in significant decrease in the neutrophils population relative to the isotype group (Figure 6B). However, we did not find total depletion or basal levels of neutrophils in the anti-Ly6G antibody therapy group **(Figure 6B)**. Additionally, 1A8 antibody treatment resulted in a significant percentage decrease (∼1%) in the monocytes population with no significant changes in the macrophages **(Figure 6B)**. Although, treatment with 1A8 antibody resulted in a significant percentage reduction (5%) in the population of CD4^+^ and CD8^+^ T cells in the lungs and spleen harvested from antibody-treated *vapC12* mutant mice as compared to WT **(Figure 6C)**. The reduction in CD4^+^ and CD8^+^ T cells could be due to the IL-4Ra/STAT6 mediated MDSCs activation resulting in T-cells immunosuppression. We found that treatment with 1A8 antibody did not affect overall lung and spleen bacillary burden during acute infection in the *vapC12* mutant treated with 1A8 antibody as compared to the isotype control treated mice **(Figure 6D)**. These data collectively suggest that the Δ*vapC12* induced hypervirulence phenotype is independent of the enhanced MDSCs population observed in the lungs of Δ*vapC12* infected mice.

**Figure 6:**
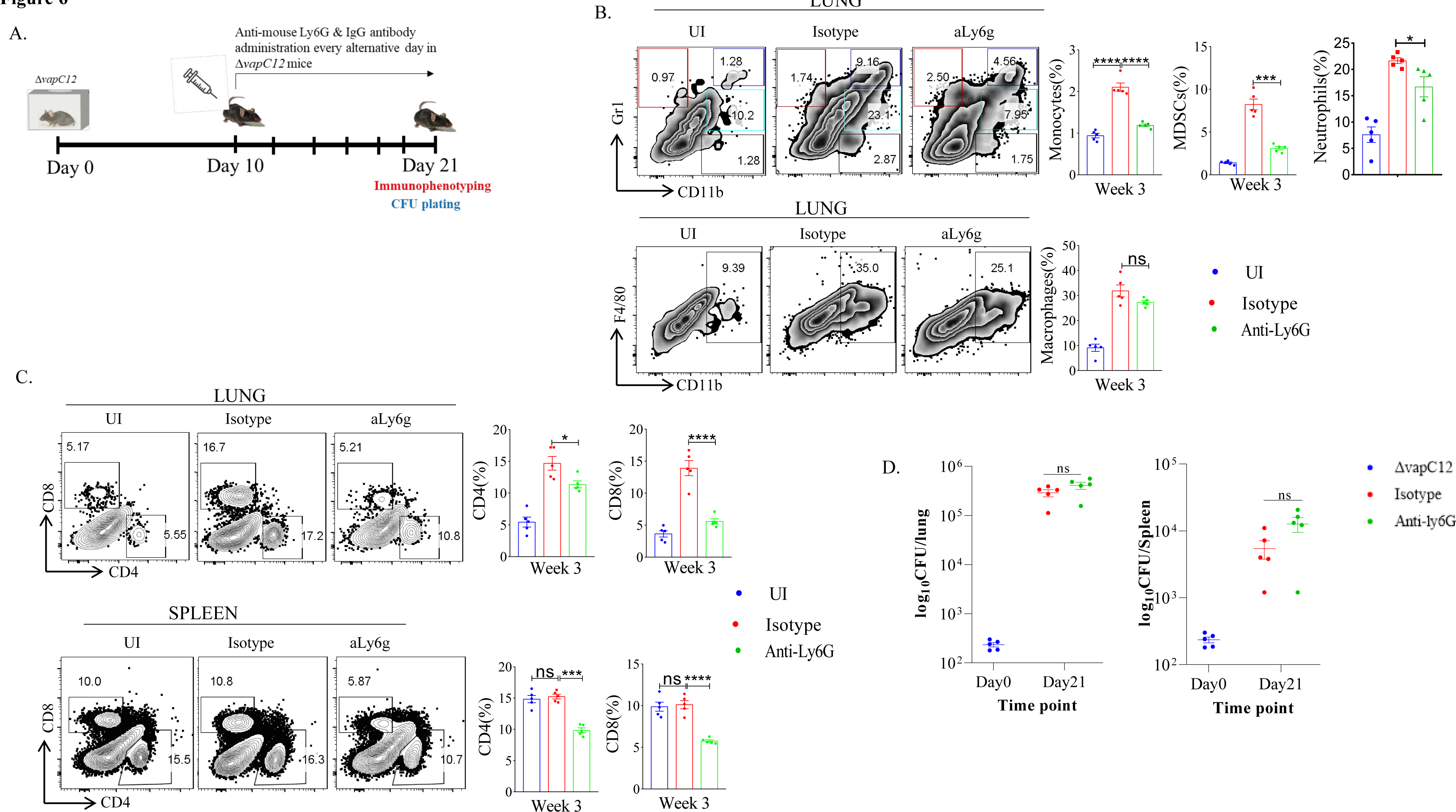
Myeloid-derived suppressor cells depletion failed to restore the *vapC12* mutant phenotype to the wild type levels. (A) Schematic representation of antibody depletion experiment in mice. *ΔvapC12* infected mice (n=5) were either treated with MDSCs depleting Anti-Ly6G or IgG control antibody every alternative day from day 10 to day 21 post-infection. Mice were sacrificed on day 21 for immunophenotyping & bacterial enumeration. (B) Zebra plots and its representative bar graph shows the percentage frequency of monocytes (Gr1+), macrophages (CD11b+F4/80+), MDSCs (CD11b+ Gr1+) and neutrophils (CD11b+Gr-1^int^) cells in the lungs. Data are shown as mean ± SEM (n=5 animals per group). Data were analyzed using the one-way ANOVA followed by Tukey’s multiple comparison test (*P <0.05, **P < 0.01, ***P<0.0005, ****P<0.0001). (C) Zebra plots and its representative bar graphs indicating percentage frequency of CD4^+^ and CD8^+^ T cells in the lung and spleen. Data were analyzed using FlowJo and shown as mean ± SEM (n=5 animals per group). Data were analyzed using the one-way ANOVA test followed by Tukey’s multiple comparison test (*P <0.05, **P < 0.01, ***p<0.0005, ****p<0.0001). (D) Scatter plot showing bacterial load in the lungs and spleen of mice infected with *ΔvapC12* at designated time points, the lungs were homogenized in 2 ml of saline, and ten-fold serial dilutions of homogenates were plated on 7H11+OADC plates. Each group constituted five mice per time point. Data plotted represent the mean ± SEM. Significant differences observed between groups are indicated. Data were analyzed using the Mann–Whitney test with **P < 0.01 and *P <0.05

### *vapC12* deletion induced hypervirulence in *M. tuberculosis* is dependent on TLR4

Neutrophil associated proteins, S100A8/A9 function as DAMPs (danger-associated molecular pattern) and can act as a ligand of TLR4 thus activating the innate immune responses [48]. Since RNA-sequencing data showed that both S100A8/A9 neutrophil-associated markers were elevated in the lungs of v*apC12* mutant, we hypothesized that TLR4 signaling could play an important role in Δ*vapC12* mediated hypervirulent phenotype. To study the role of TLR4 signaling, we infected C57BL/6 and C3H/HeJ mice (carrying mutated TLR4) with both the WT and Δ*vapC12* strains **(Figure 7A)**. As expected, relative to the WT, we observed an increase in the bacterial load in the organs isolated from C57BL/6 mice infected with the Δ*vapC12* strain. Surprisingly, the above phenotype observed in the C57BL/6 mice was completely reversed in organs isolated from C3H/HeJ mice infected with Δ*vapC12* **(Figure 7B &C).** We observed a ∼10-fold & ∼40-fold reduction in bacterial growth in lungs & spleen respectively isolated from C3H/HeJ mice infected with Δ*vapC12* as compared to C57BL/6 **(Figure 7B &C).** Further, we found a significant reduction in the population of MDSCs in the organs isolated from Δ*vapC12* infected C3H/HeJ mice as compared to WT (p < 0.05). We also observed that the neutrophil and macrophage population isolated from the lungs of Δ*vapC12* infected TLR4 defective mice were also restored to the WT levels **(Figure 7D)**. Surprisingly, restoration of the adaptive immune response in Δ*vapC12* infected TLR4 defective mice were observed only in the spleen **(Figure 7E)**. We believe that spleen being a secondary lymphoid organ is resided first by the lymphocytes and that the failure of the CD4^+^ and CD8^+^ T-cell to populate the lungs would have been restored at a later time point. These findings were in line with our bacterial enumeration data suggesting that TLR4 signaling pathway could play an important role in Δ*vapC12* mediated hypervirulent phenotype observed in the mice.

**Figure 7:**
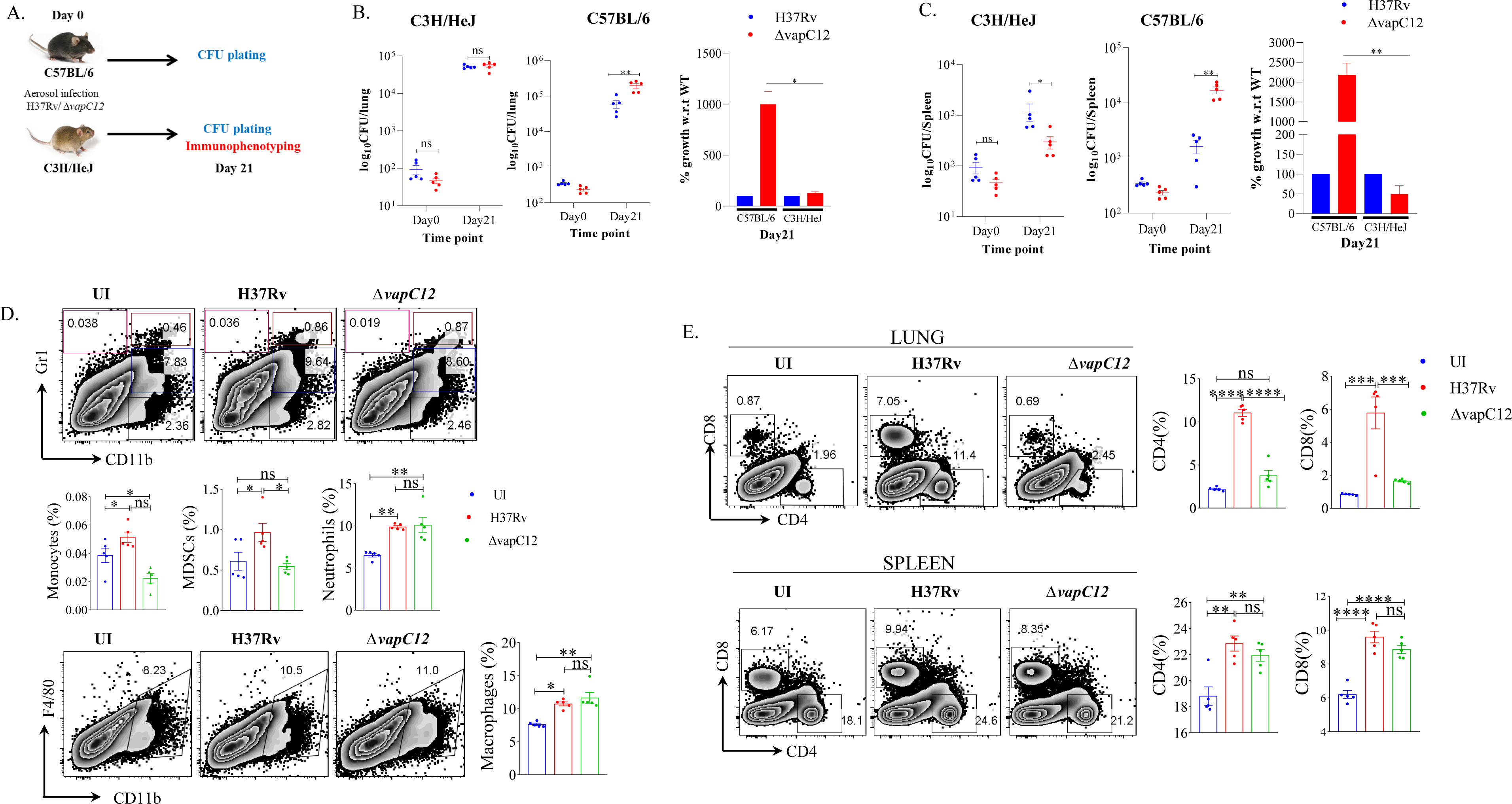
*vapC12* deletion induced hypervirulence in *M. tuberculosis* is dependent on TLR4. (A) Schematic representation of the animal experiment. C3H/HeJ and C57BL/6 mice were aerosol infected with H37Rv and *ΔvapC12*. Mice were sacrificed on indicated time point for bacterial enumeration (C3H/HeJ and C57BL/6) and immunophenotyping (C3H/HeJ). (B and C) Scatter plot showing bacterial load in the (B) lungs and (C) spleen of mice infected with H37Rv and *ΔvapC12* at indicated time points, the lungs were homogenized in 2 ml of saline, and ten-fold serial dilutions of homogenates were plated on 7H11+OADC plates. Each group constituted five mice per time point. Bar graph depicts percentage plot showing growth of *vapC12* mutant with respect to wild-type. Data plotted represent the mean ± SEM. Significant differences observed between groups are indicated. Data were analyzed using the Mann– Whitney test with **P < 0.01 and *P <0.05 (D) Zebra plot and its representative bar graphs showing the percentage frequency of monocytes (Gr1+), macrophages (CD11b+F4/80+), MDSCs (CD11b+ Gr1+) and neutrophils (CD11b+Gr-1^int^) cells in the lung. Data are shown as mean ± SEM (n=5 animals per group). Data were analyzed using the one-way ANOVA followed by Tukey’s multiple comparison test (*P <0.05, **P < 0.01, ***P<0.0005, ****P<0.0001). (E) Zebra plots and its representative bar graphs indicating the percentage frequency of CD4^+^ and CD8^+^ T cells in the lung and spleen. Data were analyzed using FlowJo and shown as mean ± SEM (n=5 animals per group). Data were analyzed using the one-way ANOVA test followed by Tukey’s multiple comparison test (*P <0.05, **P < 0.01, ***p<0.0005, ****p<0.0001).

## Discussion

Earlier we reported that the vapC12 ribonuclease toxin targets proT tRNA and modulates the levels of various proline-rich PE-PGRS proteins [22]. In the same study, we further demonstrated that organs isolated from guinea pigs infected with *M. tuberculosis* strain lacking *vapC12* gene had higher bacterial load and were found to be more pathogenic measured in terms of the degree of inflammation and pathological lesions as compared to the WT strain [22]. The current study was undertaken to further our understanding of the hypothesis that *vapC12* mediated differential expression of antigenic proteins modulates host immune response. Due to easy availability of good genetic and immunological toolbox we in the current study have decided to use mice as a surrogate host. Similar to the phenotype observed in guinea we found that the *vapC12* deficient strain of *M. tuberculosis* demonstrated a hypervirulent phenotype marked by an increase in bacterial burden and enhanced pathological lesions. Here, we have demonstrated that Δ*vapC12* induced dysregulated neutrophils influx resulting in robust transcription of S100A8/A9 proteins providing enhanced growth benefit to the mutant strain as compared to the WT. By inhibiting the S100A8/9-TLR4 axis we were able to mitigate the bacterial growth in *vapC12* mutant by reducing the detrimental inflammatory immune response.

The killing of pathogens by macrophages is known to be the primary defence mechanism against intracellular pathogens. Like T-lymphocytes, macrophages also differentiate into M1 and M2 phenotypes depending on their microenvironment and amino acid metabolic shift [37]. Arginine catabolism via iNOS leads to the production of L-citrulline and nitric oxide (NO), playing a predominant role in providing resistance against parasitic and bacterial infections. On the contrary, use of arginine as a substrate by Arg1 results in the generation of L-ornithine and urea and is known to promote disease activity [37]. During *M. tuberculosis* infection, Arg1 stimulation *in-vitro* in macrophages through a TLR-dependent and STAT6-independent pathway leads to the inhibition of NO production. Conversely, lack of Arg1 was connected to elevated NO expression and improved control of mycobacterial infection [49]. Arginine metabolism has been linked to infection as high arginase activity has been reported in lesions of leishmaniasis patients and human TB granulomas [50–52]. Most of the available piece of evidence suggests that pathogens responsible for acute as well as chronic infections manipulate STAT6-PPARγ/δ pathways to avoid M1 polarization [53–57]. While the WT *M. tuberculosis* induced polarization of macrophage towards M1, a skewed polarization towards M2 macrophages observed in mice infected with *vapC12* deficient *M. tuberculosis* strain suggests that a *vapC12*-dependent protein expression actively influences the macrophage polarization in the host. As a consequence, there is an altered T-cell response activated in terms of Th1/Th2 by the host. This implies that *M. tuberculosis* actively modulates host immunity by differentially expressing its proteins and the resulting spatiotemporal expression of *M. tuberculosis* antigens critically regulate the host immune pathways which ultimately affects the disease progression and outcome. Contrary to the WT, failure of Δ*vapC12* to prevent the accumulation of excessive neutrophils at the site of infection results in enhanced inflammation, M2 polarization and a Th2 type response which together creates an environment conducive for the growth of *M. tuberculosis* inside the host. In agreement with our observations, several reports have established that excessive neutrophil infiltration leads to tissue damage and exacerbation of inflammation which is detrimental to the host [58–61]. Additionally, upregulation of the IL-1 cytokine signaling genes suggest increased levels of proinflammatory IL-1α and IL-1β cytokines in the lungs infected with *vapC12* deficient *M. tuberculosis* strain. The dysregulation of the levels of IL-1 and IL-1 receptor antagonists cannot be ruled out.

Cathepsins carry out efficient innate and adaptive antibacterial functions, either directly by taking part in the death of invasive bacteria or by processing and presenting the antigen of infected bacteria to lymphocytes, which escalates the removal of the pathogen and triggers the development of long-lasting immunity [62]. However, bacteria have the ability to manipulate cathepsin expression and proteolytic activity in order to benefit their own intracellular survival in macrophages and escape the host-mediated immune surveillance [62]. *M. tuberculosis* avoids contact with cathepsins by modulating phagosome maturation, impairing the delivery of lysosomes to phagosomes in NF-κB dependent manner and by escaping from the phagosomes to the cytosol, thus facilitating its replication and survival [63–65]. Cystatins, which are the inhibitors of cathepsins; the proteolytic enzymes involved in the control of *M. tuberculosis* infection were increased at the transcript level in the lungs of Δ*vapC12* infected mice [43]. In support of increased peptidase inhibitor activity, we also observed suppressed levels of peptidases in lung homogenates of Δ*vapC12* infected mice. Our findings are also in concordance with previous reports that cathepsins play a central role in the immune system and hence its necessary to regulate its activity to maintain homeostasis within cells or tissues [66]. In recent years, MDSCs which are a heterogeneous population of immunocytes from myeloid origin have also been found to be a safe harbor for *M. tuberculosis* proliferation [44, 46]. As reported in recent studies, we have identified that the increase in MDSCs population mirrored alleviating protective T-cell response in the spleen of mice infected with the *vapC12* mutant as compared to the WT strain infected mice [30, 67, 68]. Depletion by anti-Ly6G antibody resulted in marked decrease in the MDSCs population and not the neutrophils in the lungs of mice infected with *vapC12* mutant strain. This inefficient depletion of neutrophils could possibly be dosage, genetic background and organ dependent. A study has reported better neutrophil depletion efficiency in naïve BALB/C mice as compared to naïve C57BL/6 mice owing to lesser neutrophils number in different organs and reduced expression of Ly6G on neutrophils in the later mice strain [69]. These results implicate that the influx of neutrophils along with MDSCs, IL-1 cytokines and endopeptidase inhibitors might be responsible for increased bacillary load and lung pathology in the mice infected with Δ*vapC12* mutant.

S100A8/A9 proteins, members of the S100 family promote the recruitment of myeloid cells, specifically neutrophils, dendritic cells and monocytes to the lungs through the induction of inflammatory cytokines [39]. Indeed, our study revealed that S100A8/A9 levels corroborated neutrophil accumulation in the lungs of mice infected with *vapC12* strain during TB disease progression. These findings are consistent with the literature available on the function of S100A8/A9 in host defense mechanisms against bacterial and viral infections [39, 70]. The RNA-sequencing data also showed increased expression of the marker gene Cxcr2 suggesting increased recruitment of mature neutrophils at week 4 post-infection in the *vapC12* mutant infected mice. Therefore, it is possible that in *vapC12* mutant infected mice, the mature neutrophils that entered the lungs conveyed a huge amount of S100A8/A9 which would have further formed a positive loop by inducing its own expression. A recent study has mechanistically demonstrated that the expression of these calprotectin is needed for the upregulation of CD11b integrin on neutrophils resulting in their accumulation during chronic TB infection [71].

S100A8/A9 plays a decisive role in innate immunity by binding to receptors such as TLR4 [48] and receptor for advanced glycation end products (RAGE), which activates multiple inflammatory pathways mediated by these pattern-recognition receptors (PRRs) [48, 72, 73]. Finally, we investigated the role of TLR4 binding in inducing S100A8/A9 mediated neutrophil over-expression in *vapC12* mutant. By blocking the TLR4 signaling (C3He/HeJ mice) the bacillary burden in the lungs of *vapC12* mice became comparable to that of the WT strain. Based on these findings, we believe that the unrestricted and extreme production of S100A8/A9 could be accountable for strong activation of TLR4 signaling, which in turn stimulates the aberrant neutrophil production resulting in an imbalance of immune response in *vapC12* mutant infected mice as compared to WT strain. The exact mechanism of S100A8/A9 overexpression and TLR4 signal activation induced in *vapC12* mutant needs to be explored further. Exploring the differentially expressed secretory proteins in *vapC12* mutant as compared to the WT strain would further give us a better insight into mechanistically understanding the downstream processes.

In conclusion, we have studied how the absence of the ribonuclease toxin in *M. tuberculosis* greatly impacted the ability of the pathogen to dampen the initial burst of neutrophils to the infected organ. The uninterrupted migration of granulocytes and continuous expression of proinflammatory cytokines resulted in enhanced pathology. The resulting M2 mediated Th-2 response, generated in the mice infected with Δ*vapC12* favored the replication and growth of *M. tuberculosi*s. Overall our data suggest that VapC12 ribonuclease toxin-mediated differential expression of *M. tuberculosis* proteins help in the timely resolution of neutrophil-driven inflammation which is critical for disease persistence in TB **(Figure 8)**.

**Figure 8:**
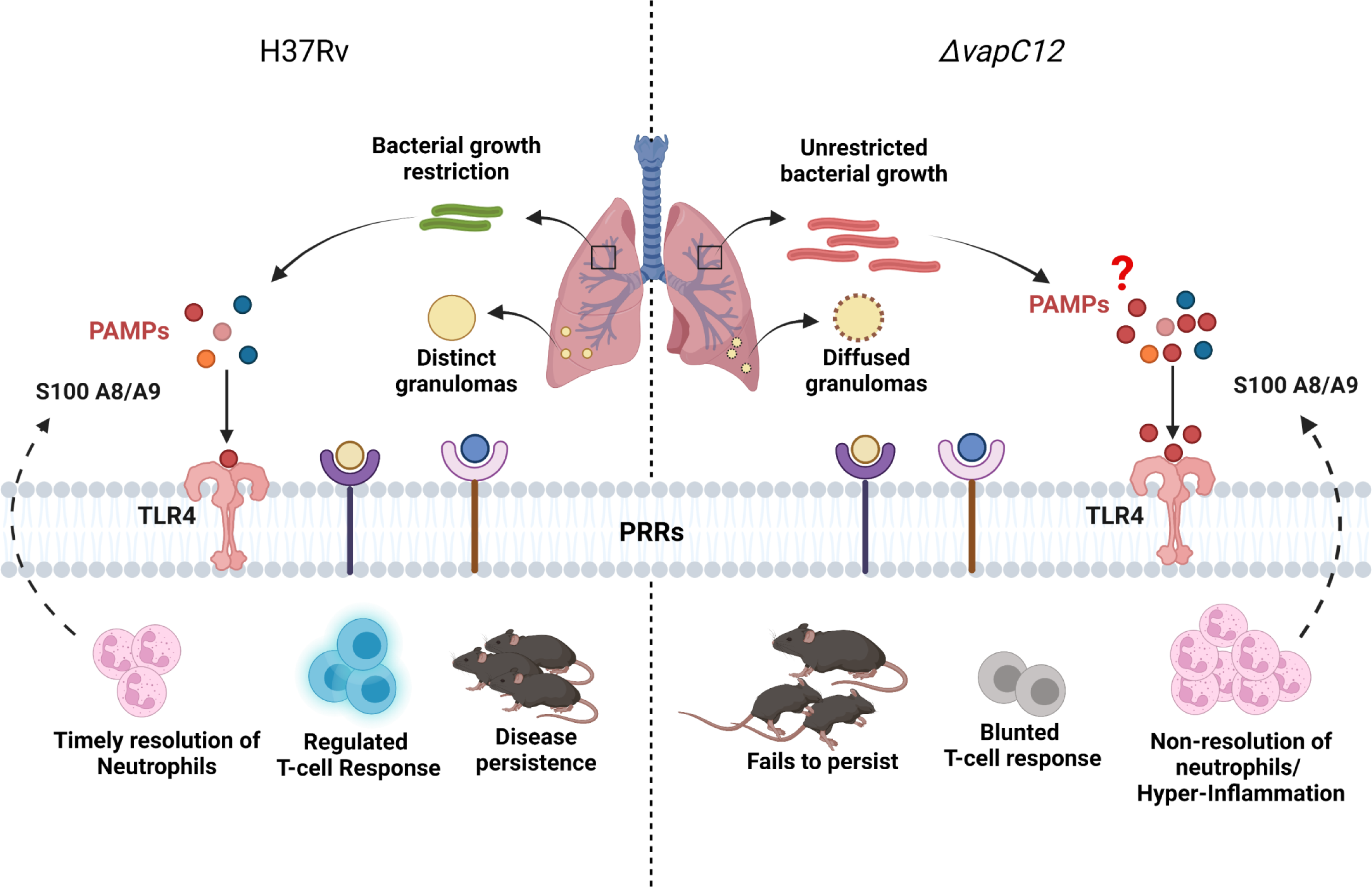
Model depicting the VapC12 toxin-mediated host immunomodulation during *M. tuberculosis* infection. Our findings demonstrate that the absence of ribonuclease toxin VapC12 favors unrestricted bacterial growth and exacerbated pathology. Additionally, *vapC12* mutant hinders the ability of *M. tuberculosis* to control the initial influx of neutrophils to the site of infection leading to continuous inflammation and blunted T-cell response. Further, non-resolution of neutrophils results in overactivation of S100A8/A9-TLR4 signaling. Finally, *vapC12* mediated expression of *M. tuberculosis* proteins is critical for disease persistence in tuberculosis.

## Statements and Declarations

### Author contributions

AP have conceptualized and supervised the study. AP, AA and ST have designed the study experiments. ST, SS, TS, MP and VN have performed the experiments. SC and AP helped in analysing the transcriptome data. SS and DR helped in analysis of flow cytometry data. ST and AKP drafted the manuscript and all authors were involved in revising it critically and have given final approval of the version to be published.

#### Acknowledgements

We are thankful to SAF and IDRF at THSTI for their support. We thank ILBS, New Delhi and histopathology facility at THSTI for histopathological analysis and Surjeet Yadav for lab maintenance. THSTI is acknowledged for providing all support for equipment and other infrastructure.

#### Data Availability

All data generated or analyzed during this study are included either in this article or in the supplementary information files. Host RNA sequencing accession number: PRJEB63988

#### Ethics approval

The animal study was reviewed and approved by the Institutional Animal Ethics Committee (IAEC), THSTI.

#### Consent for publication

Not applicable

