## Supplementary Figures for "VapC12 ribonuclease toxin modulates host immune response during *Mycobacterium tuberculosis* infection"

**Figure S1:**

A.

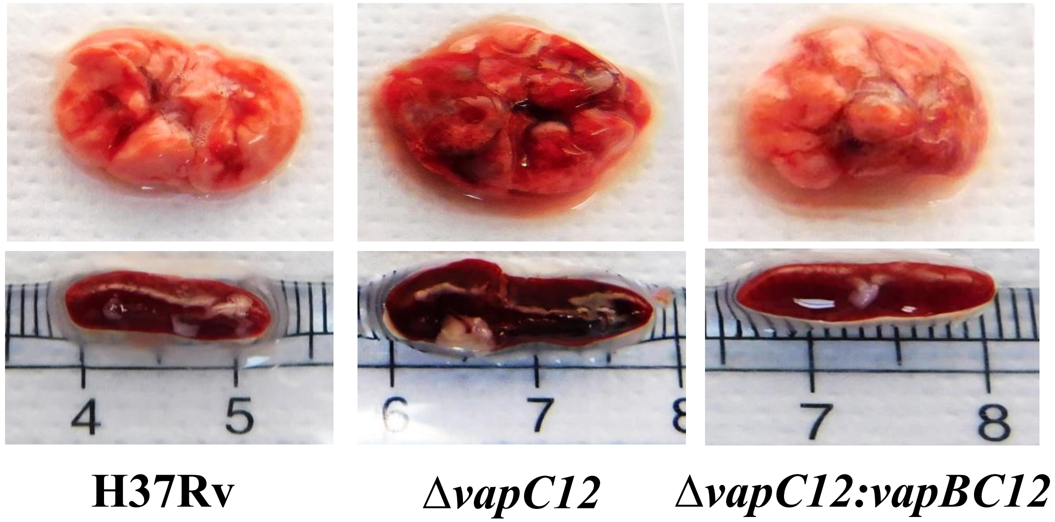

4
5
6
7
8
7
8
H37Rv
ΔvapC12
ΔvapC12:vapBC12


B.

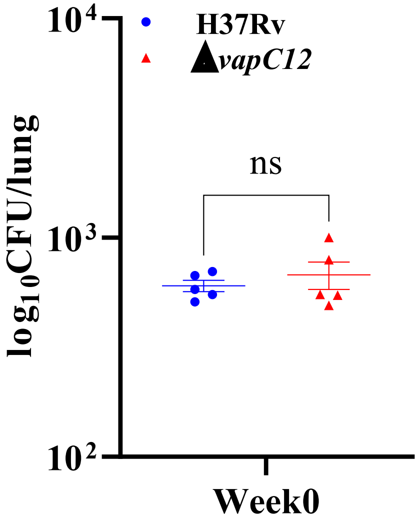

H37Rv
ΔvapC12
log10CFU/lung
ns
10^4
10^3
10^2
Week0

**Figure S2**

A.

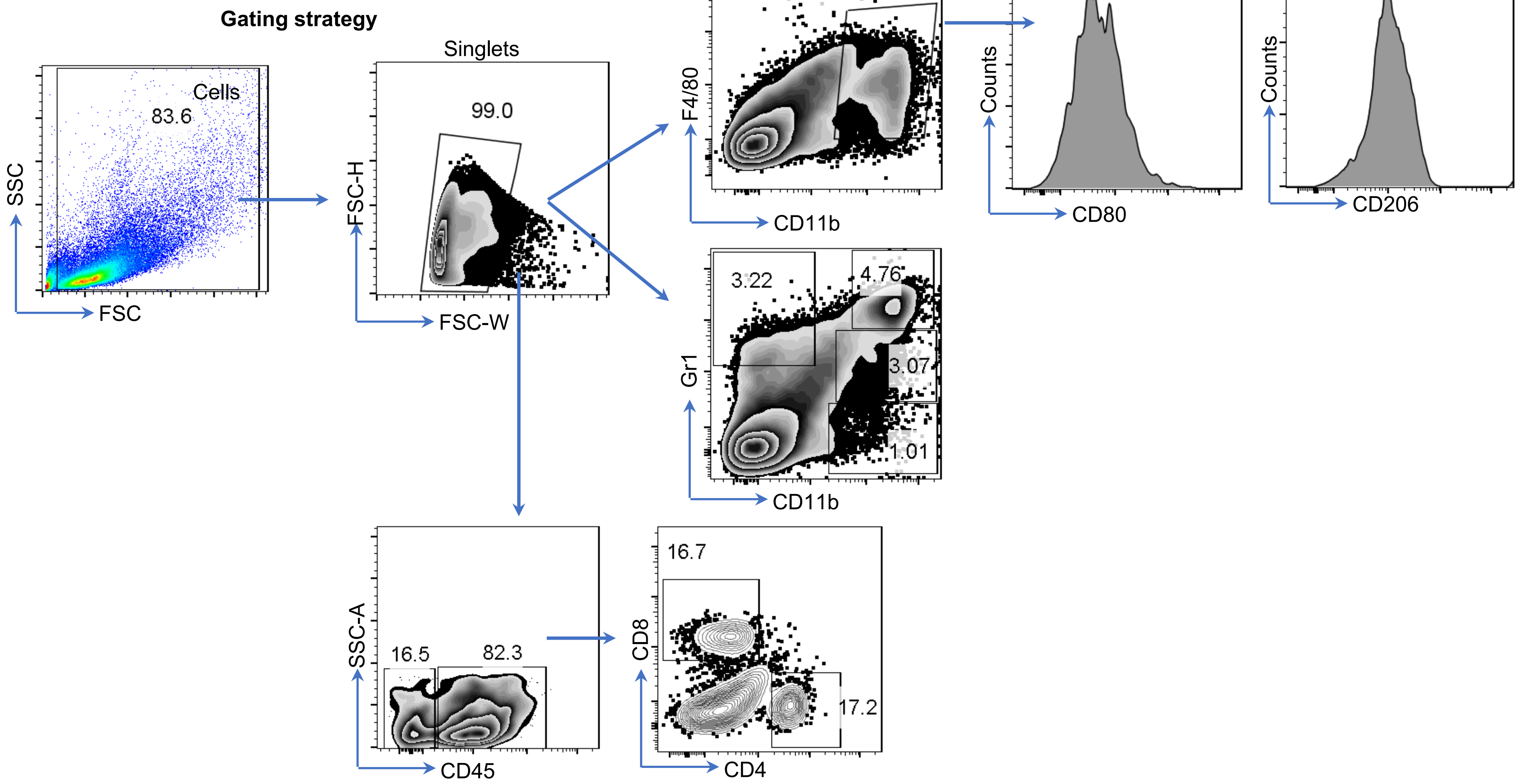
Gating strategy
Cells
83.6
SSC
FSC
Singlets
99.0
FSC-H
FSC-W
8.37
F4/80
CD11b
Counts
CD80
Counts
CD206
3.22
4.76
3.07
1.01
Gr1
CD11b
16.5
82.3
SSC-A
CD45
16.7
17.2
CD8
CD4

**Figure S3**

A.

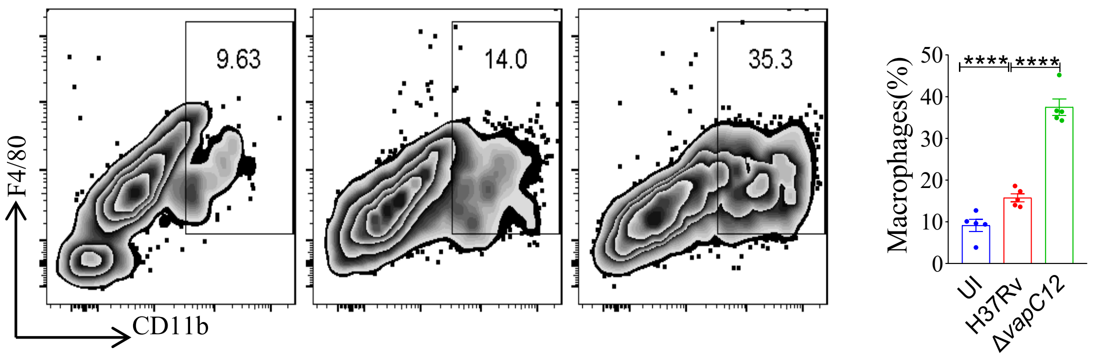

9.63
14.0
35.3
F4/80
CD11b
Macrophages(%)
****
****
50
40
30
20
10
0
UI
H37Rv
ΔvapC12

**Figure S4**

A.

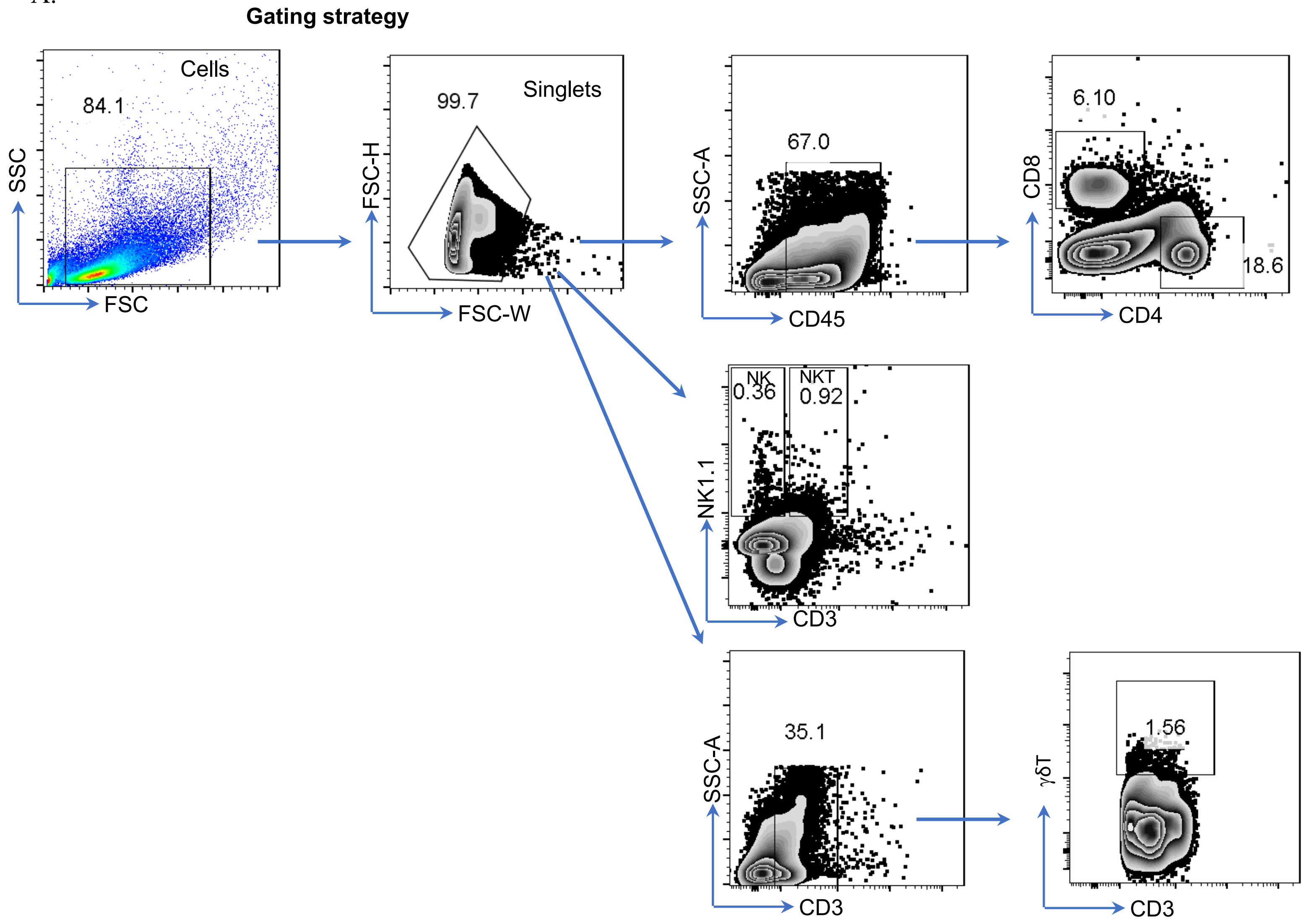

Gating strategy
Cells
84.1
SSC
FSC
99.7
Singlets
FSC-H
FSC-W
67.0
SSC-A
CD45
6.10
18.6
CD8
CD4
NK
0.36
NKT
0.92
NK1.1
CD3
35.1
SSC-A
CD3
1.56
γδT
CD3

**Figure S5**

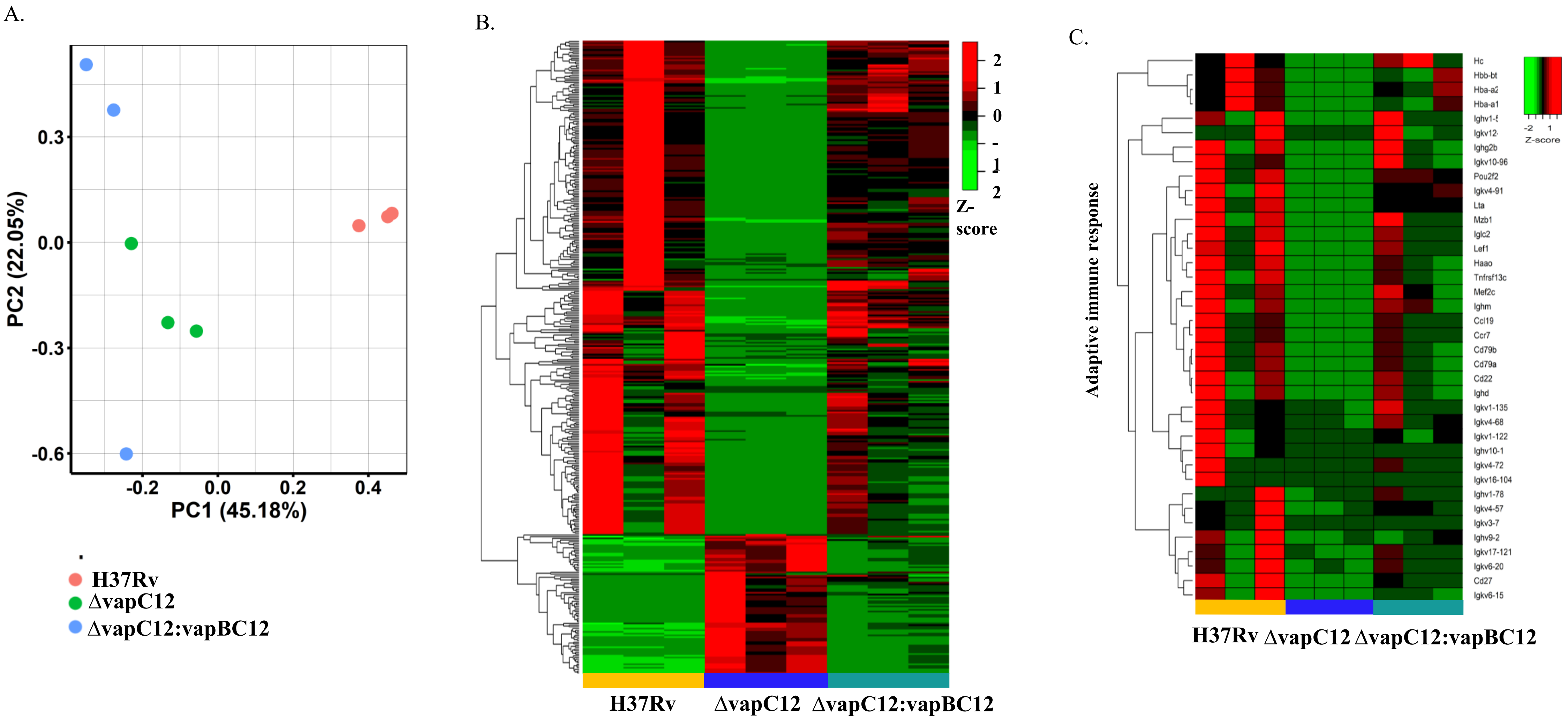
A.
0.3
0.0
-0.3
-0.6
PC2 (22.05%)
-0.2
0.0
0.2
0.4
PC1 (45.18%)
H37Rv
ΔvapC12
ΔvapC12:vapBC12
B.
2
1
0
-
1
2
Z-
score
H37Rv
ΔvapC12
ΔvapC12:vapBC12
C.
Adaptive immune response
Hc
Hbb-bt
Hba-a2
Hba-a1
Ighv1-5
Igkv12-
Ighg2b
Igkv10-96
Pou2f2
Igkv4-91
Lta
Mzb1
Iglc2
Lef1
Haao
Tnfrsf13c
Mef2c
Ighm
Ccl19
Ccr7
Cd79b
Cd79a
Cd22
Ighd
Igkv1-135
Igkv4-68
Igkv1-122
Ighv10-1
Igkv4-72
Igkv16-104
Ighv1-78
Igkv4-57
Igkv3-7
Ighv9-2
Igkv17-121
Igkv6-20
Cd27
Igkv6-15
-2
1
Z-score
H37Rv
ΔvapC12
ΔvapC12:vapBC12

### Supplementary Figures

**Figure S1:**

(A) Gross pathology of the lungs and spleen of mice infected with various strains of *M. tuberculosis* at 7 weeks post infection**.**

(B) Bacterial load enumeration using CFU to determine the initial infection by aerosol exposure in the C57Bl/6 mice lungs infected with H37Rv and *vapC12* mutant strain in the survival study.

**Figure S2**: (A) Gating strategy for Macrophages (M1 and M2 population), MDSCs, neutrophils and monocytes population.

**Figure S3:** (A) Zebra plot and its representative bar graph showing the percentage frequency of macrophages (CD11b+F4/80+) harvested from lungs. Data are shown as mean ± SEM (n=5 animals per group). Data were analyzed using the one-way ANOVA followed by Tukey's multiple comparison tests (*P <0.05, **P < 0.01, ***P<0.0005, ****P<0.0001).

**Figure S4**: (A) Gating strategy for innate ( NK, NKT and γδT ) and adaptive immune cell population ($CD4^+$ and $CD8^+$ CELLS).

**Figure S5:**

(A) Principal component analysis (PCA) of the transcriptomic data obtained from lung tissues of mice infected with WT, Δ*vapC12* and complement strain after 4 weeks of infection. The data shown are obtained from three biological replicates.

(B) Heatmap representing the expression patterns of the 372 differentially expressed genes (DEG) obtained from comparing gene expression profiles from mice infected with parental and ΔvapC12 strain of *M. tuberculosis*. Data were Z-score transformed. The data shown are obtained from three biological replicates, and the names of the corresponding strains are mentioned at the bottom of the figure. DEGs were obtained using the fold change cut-off value of 2 and FDR-adjusted p-value < 0.05.

(C) Heatmap representing the expression levels of the 38 down-regulated genes involved in the adaptive immune response. Data were Z-score transformed and the name of the genes are displayed on the right. The names of the corresponding infected strains for each sample are mentioned at the bottom of the figure.
