## Supplementary material for "VapC12 ribonuclease toxin modulates host immune response during *Mycobacterium tuberculosis* infection": Table 1

| **Primer** | **Sequence (5-3)** |
| --- | --- |
| Ifnγ-F | TGAGTATTGCCAAGTTTGAG |
| Ifnγ-R | CTTATTGGGACAATCTCTTCC |
| Tnfα-F | GGATGAGAAGTTCCCAAATG |
| Tnfα-R | TGAGAAGATGATCTGAGTGTG |
| IL-6-F | TCCTTCAGAGAGATACAGAAAC |
| IL-6-R | TTCTGTGACTCCAGCTTATC |
| IL-10-F | CAGGACTTTAAGGGTTACTTG |
| IL-10-R | ATTTTCACAGGGGAGAAATC |
| IL-2-F | TGTAAAACTAAAGGGCTCTG |
| IL-2-R | GCAGGAGGTACATAGTTATTG |
| Arg1-F | AGAAATTTACAAGACAGGGC |
| Arg1-R | ACTTAGGTGGTTTAAGGTAGTC |
| Fizz1-F | AATCCAGCTAACTATCCCTC |
| Fizz1-R | GTATCTCCACTCTGGATCTC |
| YM1-F | ACCAGGAAAGTACACAGATG |
| YM1-R | TAAATTGTTGTCCTTGAGCC |
| Cd80-F | GTAATAACAGTCGTCGTCATC |
| Cd80-R | GCCACATAATACCATGTATCC |
| Cd86-F | ACAGAGAGACTATCAACCTG |
| Cd86-R | GAATTCCAATCAGCTGAGAAC |
| Il-4-F | CTGGATTCATCGATAAGCTG |
| Il-4-R | TTTGCATGATGCTCTTTAGG |
| GAPDH-F | CGTCCCGTAGACAAAATGGT |
| GAPDH-R | TTGATGGCAACAATCTCCA |
